# Detailed phylogenetic analysis of SARS-CoV-2 reveals latent capacity to bind human ACE2 receptor

**DOI:** 10.1101/2020.06.22.165787

**Authors:** Erin Brintnell, Mehul Gupta, Dave W Anderson

## Abstract

SARS-CoV-2 is a unique event, having emerged suddenly as a highly infectious viral pathogen for human populations. Previous phylogenetic analyses show its closest known evolutionary relative to be a virus detected in bats (RaTG13), with a common assumption that SARS-CoV-2 evolved from a zoonotic ancestor via recent genetic changes (likely in the Spike protein receptor binding domain – or RBD) that enabled it to infect humans. We used detailed phylogenetic analysis, ancestral sequence reconstruction, and *in situ* molecular dynamics simulations to examine the Spike-RBD’s functional evolution, finding that the common ancestral virus with RaTG13, dating to at least 2013, possessed high binding affinity to the human ACE2 receptor. This suggests that SARS-CoV-2 likely possessed a latent capacity to bind to human cellular targets (though this may not have been sufficient for successful infection) and emphasizes the importance to expand the cataloging and monitoring of viruses circulating in both human and non-human populations.

## Introduction

Viral pathogens are a continuous and evolving challenge for human populations.^1,2^ Most known viruses maintain species-specific infectivity, often co-evolving with their host to mirror animal species trees.^3,4^ While less common, the emergence of novel viral pathogens is of particular interest because they often exhibit abnormal degrees of infectivity and/or virulence,^5^ having not evolved to a natural selection balance with their new host.^6^ Viruses of animal origin include periodic Ebola outbreaks,^7^ the 1918 “Spanish Flu”,^8^ and most recently, SARS-CoV-2, the viral agent that causes COVID-19.^9^ In these cases, viruses spread through human populations after evolving to “cross the species barrier”.^10^ Yet, many questions remain for viruses of non-human origin: How do they acquire the ability to infect humans? Is it wholly dependent on “recognition” (a function typically mediated by protein-protein binding between viral capsid and target host cell), or must there be changes in other viral replication mechanisms as well? And specifically focusing on SARS-Cov-2, did it evolve to infect humans via many evolutionary changes or only a few? Was it dependent on a key functional shift in its ability to bind human cells, or is there evidence that other genomic changes were needed for it to acquire its strikingly high degree of infectivity? Answering these questions is critical if we are to understand both the origin of specific viruses, such as SARS-CoV-2, as well as the capacity of animal viruses to evolve human infectivity.

SARS-CoV-2 emerged in late 2019^11^ and has high infectivity, spreading rapidly around the world, causing a global health emergency.^12^ A member of the Coronaviridae family of polymorphic, enveloped, single stranded RNA viruses,^13^ it is thought that SARS-CoV-2 evolved from a zoonotic origin,^14,15^ owing to its clear evolutionary relationship with coronaviruses that have been isolated from animals^16^ (its closest known evolutionary relative is the bat coronavirus, RaTG13^17–20^ and the second-closest known relative is a pangolin coronavirus, Pangolin-CoV).^21^ While most of the SARS-CoV-2 genome mirrors the RaTG13 genome, some genomic regions, including the Spike glycoprotein Receptor Binding Domain (RBD) (which mediates viral entry into host cells), have greater sequence similarity to the Pangolin-CoV homolog,^22^ prompting some to suggest SARS-CoV-2 may be the product of recombination during co-infection.^21–24^

The Spike protein is a key component of the SARS-CoV-2 infection pathway.^25^ Knockout and overexpression studies have demonstrated that binding of the Spike-RBD to human angiotensin converting enzyme 2 (hACE2) mediates cellular entry of SARS-CoV-2.^26–30^ The SARS-CoV-2 Spike protein binds the hACE2 receptor with greater affinity than the SARS-CoV-1 homolog, suggesting as a possible explanation for its greater infectivity.^29^ As a result, it has been suggested that high affinity for hACE2 is a recently acquired trait for the SARS-CoV-2 Spike-RBD.^32–34^ Given this, a critical question remains to be answered: How and when did the SARS-CoV-2 Spike protein evolve its relatively higher affinity for the hACE2?

With this question in mind, we set out to robustly characterize the evolutionary changes that accompanied the emergence of SARS-CoV-2, and that distinguish it from its closest zoonotic relatives. We focused on the evolution of the Spike-RBD by leveraging its known evolutionary relationships to other animal and human viruses and employed ancestral sequence reconstruction in conjunction with molecular dynamics simulations to identify biochemical and biophysical changes that enhanced Spike binding to the hACE2 receptor.

## Results/Discussion

Our initial phylogenetic analysis, performed to provide context, utilized whole viral genomic data, and generally supports prior studies’ conclusions, finding similar levels of nucleotide identity to the RaTG13 genome (96.0% sequence identity) and the Guangxi Pangolin-CoV genome (90.0% sequence identity) (**Supplementary Figure 1**).^21,29^ Next, we sought to investigate the evolution of SARS-CoV-2 infectivity by performing ancestral sequence reconstruction for the Spike-RBD amino acid sequence (**Figure 1A**). While cross-species protein sequence comparisons have been used to investigate critical amino acid changes in the SARS-CoV-2 Spike protein,^37^ leveraging the phylogenetic relationships allows us to infer the most likely ancestral sequences for this protein allows us to focus on the subset of genetic changes that are specific to its recent evolution.^38^ We inferred statistically well-supported reconstructions of the Spike-RBD sequence for both the common ancestor of all human SARS-CoV-2 cases (labelled “N1”, **Figure 1A,C**) and the its common ancestor with the closest animal virus (labelled “N0”, **Figure 1A,C**). N1 is identical to the Spike-RBD in the SARS-CoV-2 reference sequence, as expected, while the N0 Spike-RBD sequence is, to our knowledge, unique, reflecting the uniqueness of SARS-CoV-2’s viral origin.^21,39^ N0 differs from N1 at 4 positions (346, 372, 498, and 519 – **Figure 1B**). The reconstruction of N1 for each of those positions is statistically well-supported, with a posterior probability (P.P.) of 1 obtained from two independent calculations (**Supplemental Table 1; Supplementary Methods**). The reconstruction for N0 has high statistical support for positions 346, 372, and 519 (P.P. > 0.94), while position 498 was ambiguously reconstructed, with two alternate states comparably probable (**Supplemental Table 1**). All other positions were reconstructed with high confidence (P.P.>0.85). Together, these four changes (t346R, t372A, h/y498Q, and n519H) differentiate the evolved SARS-Cov-2 Spike protein from the most recent common ancestor with animal viruses (**Figure 1**). As such, this ancestral virus must have existed at least as early as 2013 (as one of its descendants – RaTG13 – was isolated in that year), meaning that the branch between the N0 and N1 ancestors covers at least 7 years of molecular evolution (**Figure 1A**).

**Figure 1:**
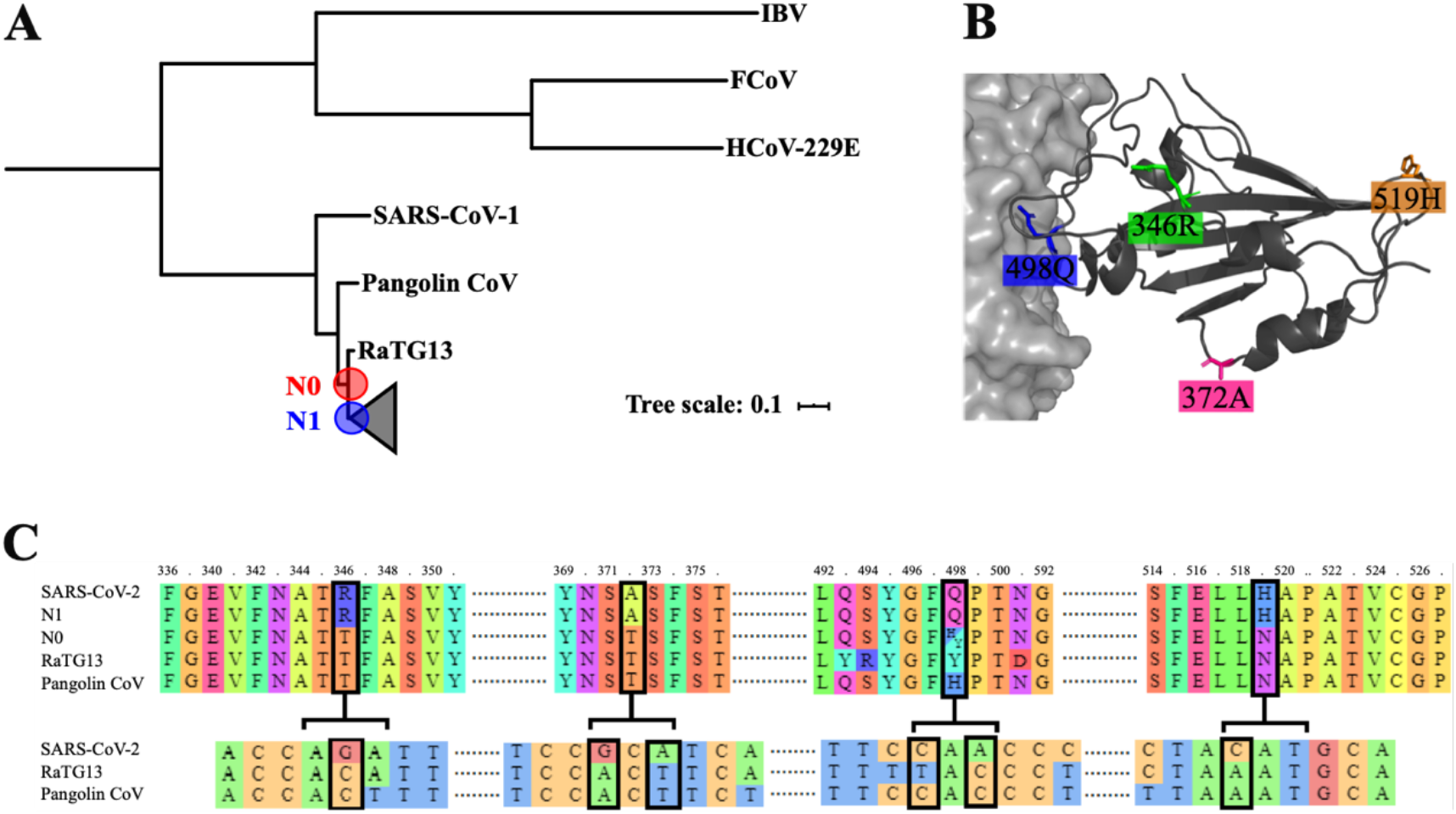
Detailed examination of SARS-CoV-2 evolution. **A**. Phylogeny illustrating the last common ancestor all SARS-CoV-2 Spike-RBDs (N1) and of SARS-CoV-2 and the RaTG13 Spike-RBD (N0). **B**. Structural representation of the four mutations in the Spike-RBD (ribbon diagram) relative to the ACE2 receptor (Space filling model) that differ between N0 to N1. Stick models show the mutations in their N1 state. **C**. Alignment of the of the Spike-RBD of SARS-CoV-2 and its ancestors for both protein (top) and DNA (bottom). Black boxes highlight the four mutations that differ from N0 to N1.

To quantify functional differences between the N0 ancestor and the Spike-RBD sequences, we conducted 10 ns molecular dynamics simulations (see **Supplementary Methods**) of the Spike Receptor Binding Domain (RBD) in complex with hACE2 (starting point for each simulation was modelled off crystal structures of the SARS-CoV-2 Spike-RBD/hACE2 complex),^27^ which we used to calculate the free energy contributions from electrostatics, polar solvation, van Der Waals interactions, and solvent-accessible surface area (SASA) to infer the free energy of binding between those two proteins.^40,41^ We quantified the root-mean-squared deviation (RMSD) of the portion of the RBD closest to the hACE2 receptor (residues 397 to 512) for each of our replicates to confirm complex stability (**Supplementary Figure 2**). Contrary to our expectations, the free energy of binding between the Spike-RBD and the hACE2 appears to have decreased between N0 and N1. In fact, each of the 4 changes either reduced or did not significantly change the free energy of binding (**Figure 2A**) (this is true for both alternate reconstructions of position 498 in N0). While overall these four historical changes reduced binding affinity to hACE2, they did not do so equally: t346R and h/y498Q showed the largest decreases in affinity (**Figure 2B**). Additionally, there was no evidence of an epistatic relationship among the four substitutions (see **Supplementary Methods**).

**Figure 2:**
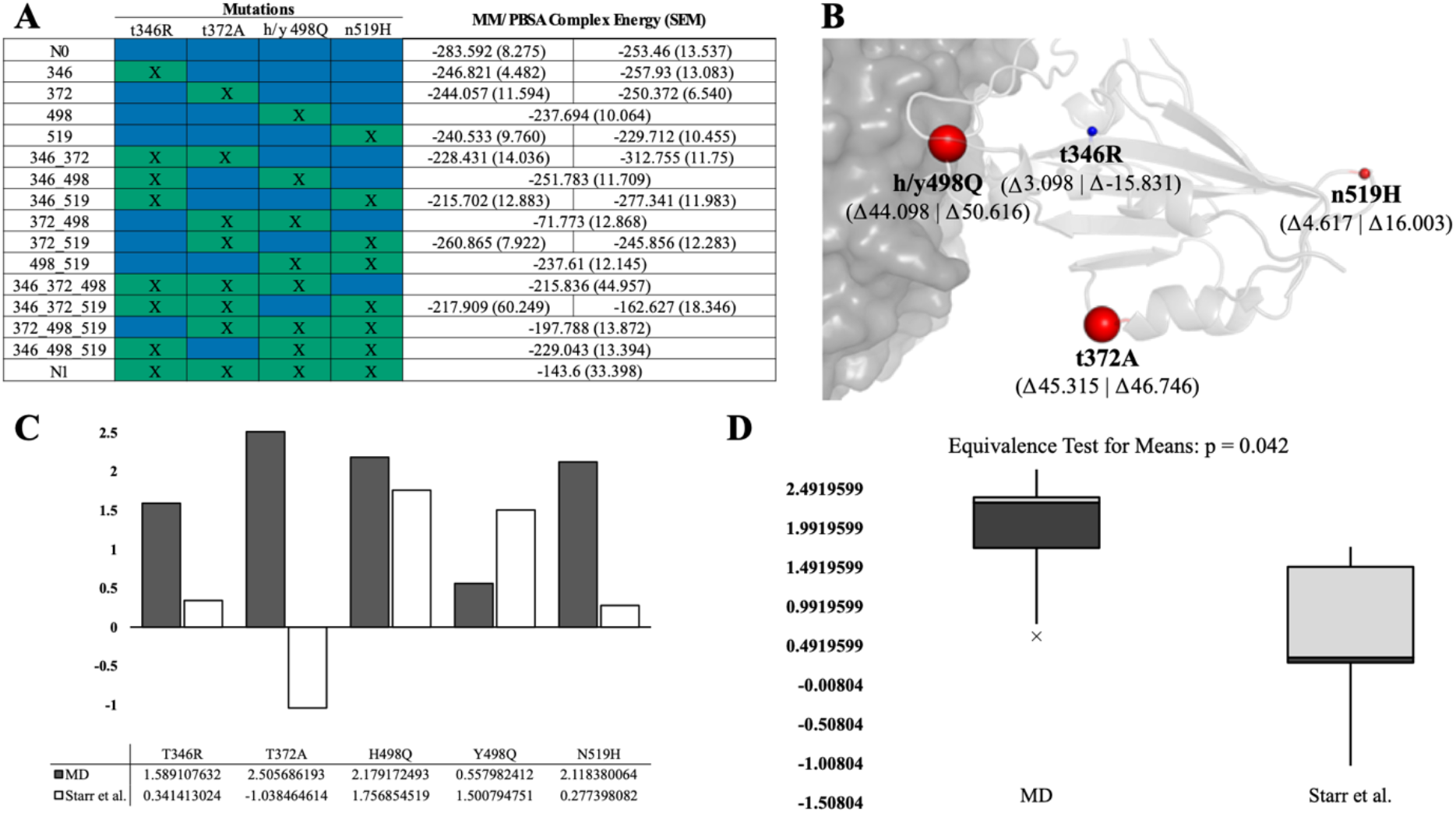
Characterization of SARS-CoV-2 Spike-RBD functional effects of evolution. A. Table of MM/PBSA binding energies between receptor binding domains of SARS-CoV-2 evolutionary constructs and hACE2 receptor (note that lower energy indicates tighter binding). Blue cells indicate the presence of the ancestral (N0) state and green cells (with an “x”) indicate the presence of the SARS-CoV-2 state (N1) at a given position. Two values are present for constructs with an ancestral (N0) state at position 498 (which reflect the ambiguity of its ancestral reconstruction), corresponding to h498 and y498 from left to right. Energies are shown as the mean of three replicate simulations with SEM indicated in parenthesis. B. Relative effect of changes in SARS-CoV-2 receptor binding domain from ancestral (N0) to SARS-CoV-2 (N1) state on MM/PBSA binding energies. Size of spheres indicate the relative magnitude, with red spheres indicating decreased binding affinity and blue indicating increased binding affinity. Values are averaged for h498 and y498 states (both raw values shown in parentheses). C. Comparison of molecular dynamics and *in vitro* z-score normalized changes in binding energy for each mutation from N0 to N1. Changes are shown relative to the z-score normalized current (N1) binding energy. D. Box plot illustrating the range of z-score normalized changes in binding energy for the *in vitro* and molecular dynamics data. An equivalence test for means (p = 0.042) suggests that the molecular dynamics simulation reflects *in vitro* data.

Somewhat surprised by these reductions in binding energy between the N0 and N1 state, we set out to confirm that our observations were not an outcome of oversimplifications in the MD forcefield. We compared standardized changes in binding energy in our MD data and recently released *in vitro* data ^42^ and saw remarkably similar changes in binding energy (p = 0.042, Equivalence Test for Means) (**Supplemental Table 3**). In particular, both alternative reconstructed states for position 498 in N0 clearly improve hACE2 binding affinity in both our simulations and in *in vitro* functional measurements (**Figure 2; Supplemental Table 3**).^42^ There was a discrepancy between the two datasets at position 372 and future *in vitro* experiments are required to determine the combined effects of this mutation with the other substitutions (**Figure 2; Supplemental Table 3**).

Nonetheless, these results demonstrate that, contrary to expectations, evolutionary changes since 2013 did not improve the SARS-CoV-2 Spike-RBD’s binding with hACE2. While there are other animal coronaviruses known to bind to the hACE2 receptor,^43^ to our knowledge, this is the first demonstration that the SARS-CoV-2’s common ancestor with the RaTG13 lineage may have been capable of binding to hACE2. This has important implications for our understanding of how SARS-CoV-2 evolved to infect humans. First, it suggests that the binding affinity between the Spike-RBD and hACE2 may not be a critical driver of the high degree of infectivity that has been observed during its recent outbreak. Instead, it suggests that tight hACE2 binding may be a latent property of this virus, and that high infectivity may instead have emerged via a distinct set of molecular changes in the SARS-CoV-2 genome. Second, it calls into question the presumption of a recent zoonotic origin for this disease; while other molecular components of the current SARS-CoV-2 virus may have acquired recent evolutionary changes that promoted its infectivity in humans, it appears that the high affinity for hACE2 was not among them.

If this is the case – that this viral lineage possessed the ability to bind hACE2 with high affinity for at least the past 7 years (**Figure 1B**) – then why did it not emerge as a public health issue until recently? One possibility is that binding hACE2 by the Spike-RBD is not sufficient, on its own, to infect humans, and that other molecular components first needed to acquire new functions to do so. A second possibility is that this virus may have been capable of infecting human cells for a longer period of time in the past, but that its ancestral form either presented with far fewer symptoms (making it less disruptive and/or noticeable to those infected), or that it was far less infectious (thereby impacting only a small number of people), in either case escaping the notice of public health monitoring (**Figure 3**). To test this, a broad and concerted effort to sequence the range of coronaviruses across human populations would need to be conducted, in order to test whether a closely related virus may also be circulating.^44–46^

**Figure 3.**
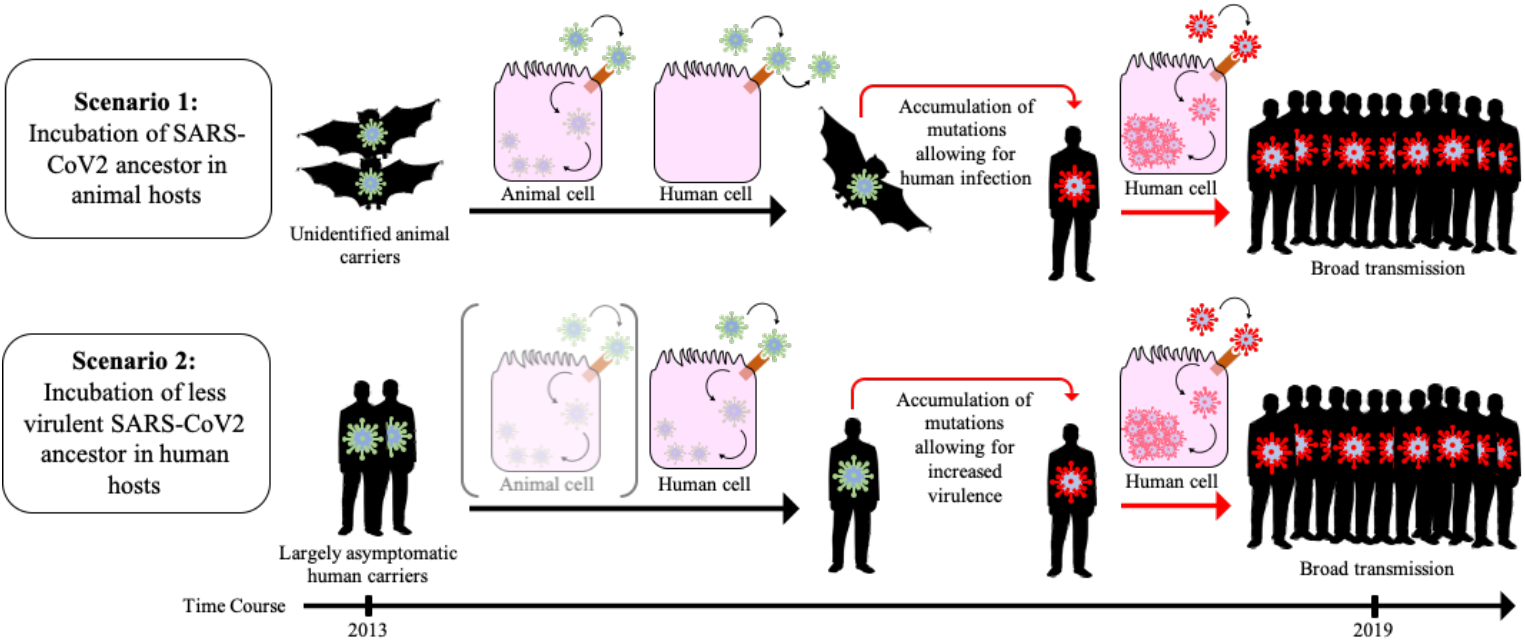
Schematic of two possible evolutionary scenarios stemming from the observed evolutionary SARS-CoV-2 Spike-RBD function. In Scenario 1, it is postulated that a zoonotic ancestral SARS-CoV-2 strain possessed the ability to effectively bind hACE2 but was unable to effectively enter human cells, requiring the presence of subsequent mutations to infect humans. In Scenario 2, an ancestral SARS-CoV-2 strain was actively infecting humans prior to the outbreak at low levels, thus escaping public health detection until subsequent mutations lead to increased infectivity and/or severity.

Naturally, as an *in silico* study, these results should be interpreted with some caution. Insofar as they can be validated, however, they are largely consistent with direct functional measurements in the lab.^42^ Ideally, combinatorial libraries could be constructed^47,48^ and functionally screened^48^ in order to glean more detailed insights into the molecular mechanisms underlying the recent evolution of this virus.

## Conclusion

Predicting the emergence of highly infectious and virulent diseases, while difficult, is vital for human population health.^50^ To do so, however, we must take steps to understand how pandemic diseases – such as SARS-CoV-2 – emerged as they did, and to understand if and when they acquired the novel molecular functions that enabled their infectivity. In this case, it appears that the SARS-CoV-2 Spike-RBD did not recently evolve binding affinity to a human-specific protein. Instead, that function appears to have been latent, making it clear that the evolution of this disease – along with so many other aspects of its etiology – is more complex than expected.

## Materials and Methods

To confirm SARS-CoV-2 etiology we constructed a phylogenetic tree using PhyML 3.0 of 26 known enzootic and endemic viruses aligned using MAFFT version 7. We then obtained the amino acid spike glycoprotein sequences from the coronaviruses in our phylogeny and 479 SARS-CoV-2 samples, by either downloading the sequences from NCBI – if available, or through extraction using nBLASTx. Spike sequences were then aligned, and ancestral sequence reconstruction was performed using two software packages (RaxML and GRASP).

Next, we performed *in silico* mutagenesis of x-ray crystallography structures for the SARS-CoV-2 spike protein RBD-hACE2 complex to create a library of mutational differences between the current SARS-CoV-2 spike RBD and our inferred ancestor. Molecular interactions for each structure in the library were then characterized through 10 ns MD simulations run in GROMACS. Binding energy between residues 397 to 512 of the SARS-CoV-2 spike RBD and the hACE2 receptor in the MD simulations were calculated using g_mmpbsa and genetic effects were measured for these energies.

Finally, MD energy outputs were compared to existing *in vitro* data using standard statistical methods.

## Supplementary Methods

### Confirmation of SARS-CoV-2 etiology

A phylogenetic analysis of 26 viral genomes was performed to confirm known SARS-CoV-2 ancestors. 24 known enzootic and endemic viruses and the SARS-CoV-2 reference genome and the Pangolin-CoV genome were downloaded from the National Center for Biotechnology Information (NCBI)^51^ and Lam et al.^21^ respectfully. Selected sequences were aligned using the Multiple Alignment using Fast Fourier Transform Version 7 (MAFFT) FFT-NS-2 algorithm. ^52,53^ MAFFT default parameters were used in our alignment, meaning gap penalties were assigned a value of 1.53. PhyML 3.0 was employed to construct a phylogeny of aligned genomes. ^54,55^ Bayes values ≥ 0.90 were considered statistically significant. The output tree was visualized using the online tool, Interactive Tree of Life (iTOL), and statistically significant clades were examined to validate current knowledge surrounding SARS-CoV-2 evolution.^56^

### Construction of spike glycoprotein ancestral sequence

nBLASTx, run using a BLOSUM 62 matrix, a gap opening penalty of 11 and a gap extension penalty of 1, was employed to extract the Spike glycoprotein from the 479 SARS-CoV-2 sequences obtained from GISAID^58,59^ selecting for one sequence per day per country from December 30, 2019 to March 25, 2020, and the Pangolin-CoV genome.^57^ Additional, Spike sequences, including the RaTG13 Spike protein, were obtained directly from NCBI.^51^ Protein sequences were initially aligned using the Multiple Sequence Alignment by Log-Expectation (MUSCLE) program.^60^ The optimal parameters for phylogenetic reconstruction analysis were taken from the best-fit evolutionary model selected using the Akaike Information Criterion (AIC) implemented in the PROTTEST3 software,^61^ and were inferred to be the Jones-Taylor-Thornton (JTT) model^62^ with gamma-distributed among-site rate variation and empirical state frequencies. Phylogeny was inferred from these alignments using the RaXML v8.2.9 software^63^ and results were visualized using FigTree v1.4.4 (https://github.com/rambaut/figtree/releases). Ancestral sequence reconstruction was performed with the FastML software^64^ and further validated independently using the Graphical Representation of Ancestral Sequence Predictions (GRASP) software.^65^ Statistical confidence in each position’s reconstructed state for each ancestor determined from posterior probability; any reconstructed positions with less than 95% posterior probability was considered ambiguous, and alternate states were also tested.

### Mutagenesis of ancestral proteins

To understand the evolutionary importance of sequence changes observed between ancestral, zoonotic, and SARS-CoV-2 spike protein sequences, *in silico* mutagenesis and binding energy studies were performed. A previously constructed x-ray crystallography structure for the complex between the receptor binding domain (RBD) of the SARS-CoV-2 spike protein and the human hACE2 receptor were obtained from RCSB (accession number 6M0J). Utilizing PyMOL mutagenesis wizard, ^66^ the four missense mutations (R346t, A372t, Q498h or Q498y, H519n) identified between the N0 and N1 sequences were introduced into the SARS-CoV-2 RBD sequence, replicating the sequence of the putative ancestral zoonotic (N0) sequence. In addition to the N1 and N0 structures, additional structures were developed in a similar fashion, selectively including each of the 4 mutations to represent all of the possible combinations that these mutations may have existed throughout evolutionary time

### Simulation of ACE2 interactions using molecular docking

Molecular interactions were characterized with molecular dynamics simulations using GROMACS, TIP3P waters and CHARM07 force-field parameters for proteins. For each condition, three replicate 10 ns simulations were run, starting from crystal structures or structural models. Historical mutations were introduced and energy-minimized before MD simulation. Each system was solvated in a cubic box with a 10 Å margin, then neutralized and brought to 150 mM ionic strength with sodium and chloride ions. This was followed by energy minimization to remove clashes, assignment of initial velocities from a Maxwell distribution, and 1 ns of solvent equilibration in which the positions of heavy protein and DNA atoms were restrained. Production runs were 50 ns, with the initial 10 ns excluded as burn-in. The trajectory time step was 2 fs, and final analyses were performed on frames taken every 12.5 ps. We used TIP3P waters and the CHARM07 FF03 parameters for proteins, as implemented in GROMACS 4.5.5.^67^ Analyses were performed using VMD 1.9.1.^68^ GROMACS output was uploaded into Visual Molecular Dynamics (VMD) for Root-Mean Squared Deviation (RMSD) Analysis using the RMSD trajectory tool (ref). After discovering large deviations in RMSD values for the full RBD, which we attributed to noise at the ends of the RBD, we isolated our analysis to residues 397 to 512 of the RBD.

### Measurement of binding energies

Next, we measured the binding energies between residues 397 to 512 and the ACE2 receptor using g_mmpbsa, a program which employs Molecular mechanics Poisson–Boltzmann surface area (MMPBSA) calculations to determine binding energy. Van der Waal forces, polar solvation energy, apolar solvation energy and SASA energy were calculated every 0.25 ns using a gridspace of 0.5 and a solute dielectric constant of 2. The output of the three replicates was amalgamated and binding energy was calculated using the bootstrap analysis (n = 2000 bootstraps) published by Kumari et al.^40,41^ We then characterize the genetic effect of each mutation (on average) and assessed whether there were any statistically significant epistatic interactions using established methods.^46,47^

### Comparison to in vitro data

*In vitro* changes in binding energy for the four mutation were obtained from Starr et al. ^42^. These data and our binding energies for the N1, 346_372_498, 346_372_519, 346_498_519 and 372_498_519 were each standardized using Z-scores. Changes in binding energy to the N1 state for each standardized score was calculated by subtracting the N1 energy from the mutant energy. An equivalence test for means (TOST) test was performed on the standardized changes in binding energy.

---

**Supplementary Figure 1:**
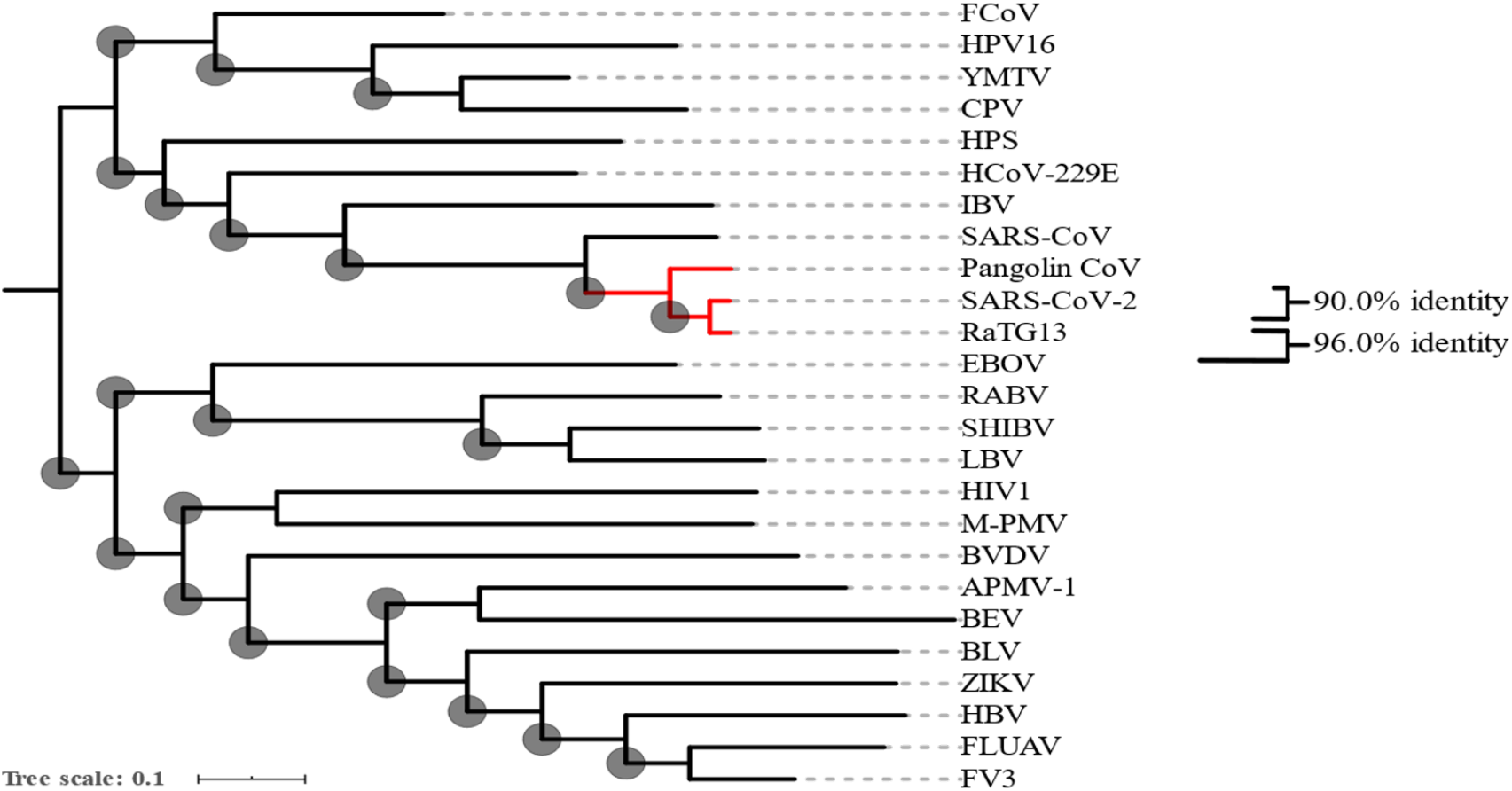
Phylogenetic reconstruction of family that includes SARS-CoV-2. Phylogeny was constructed using 25 whole genome sequencing data from a series of known coronaviruses. The two closest relatives to SARS-CoV-2 are highlighted in red and sequence identities are specified.

**Supplementary Figure 2:**
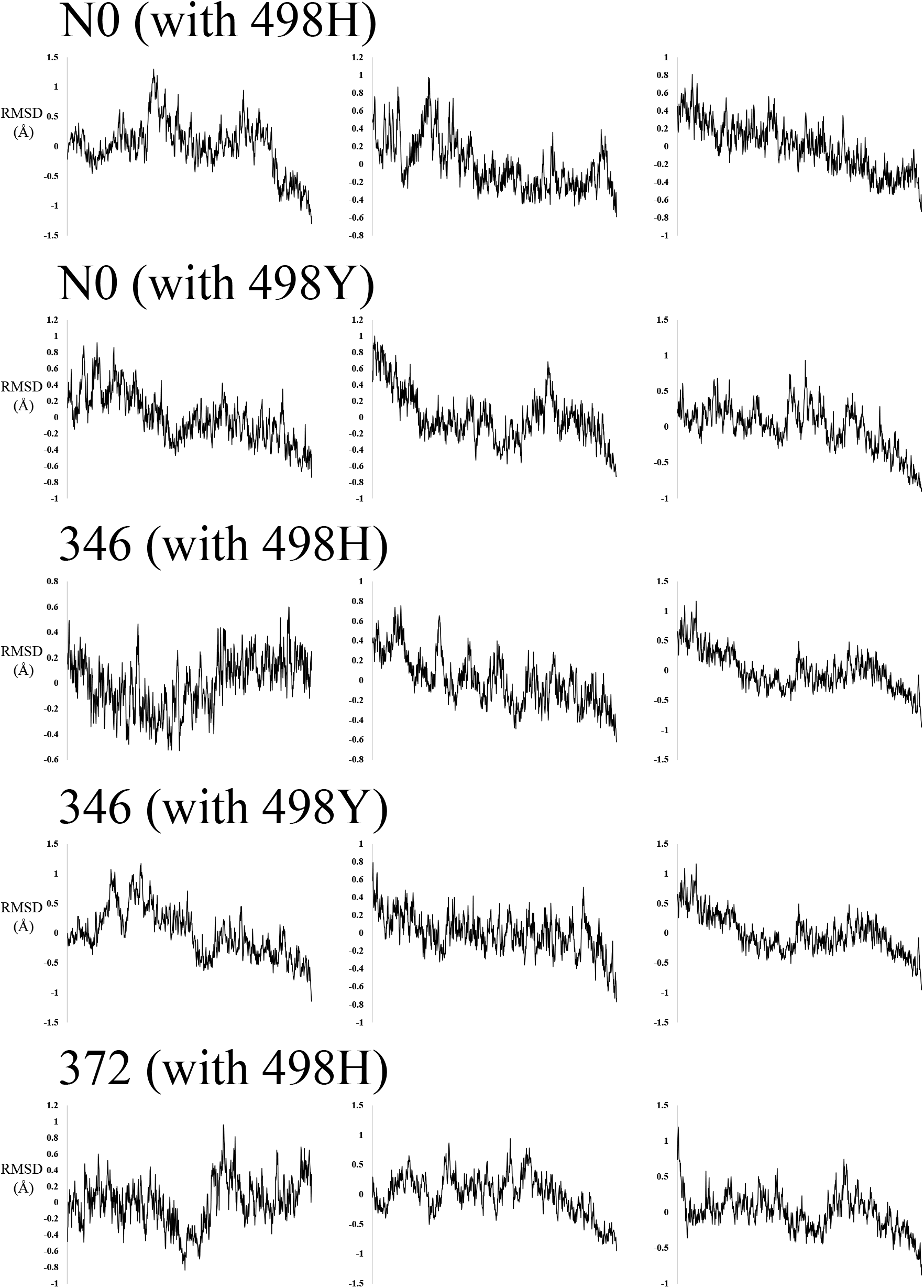
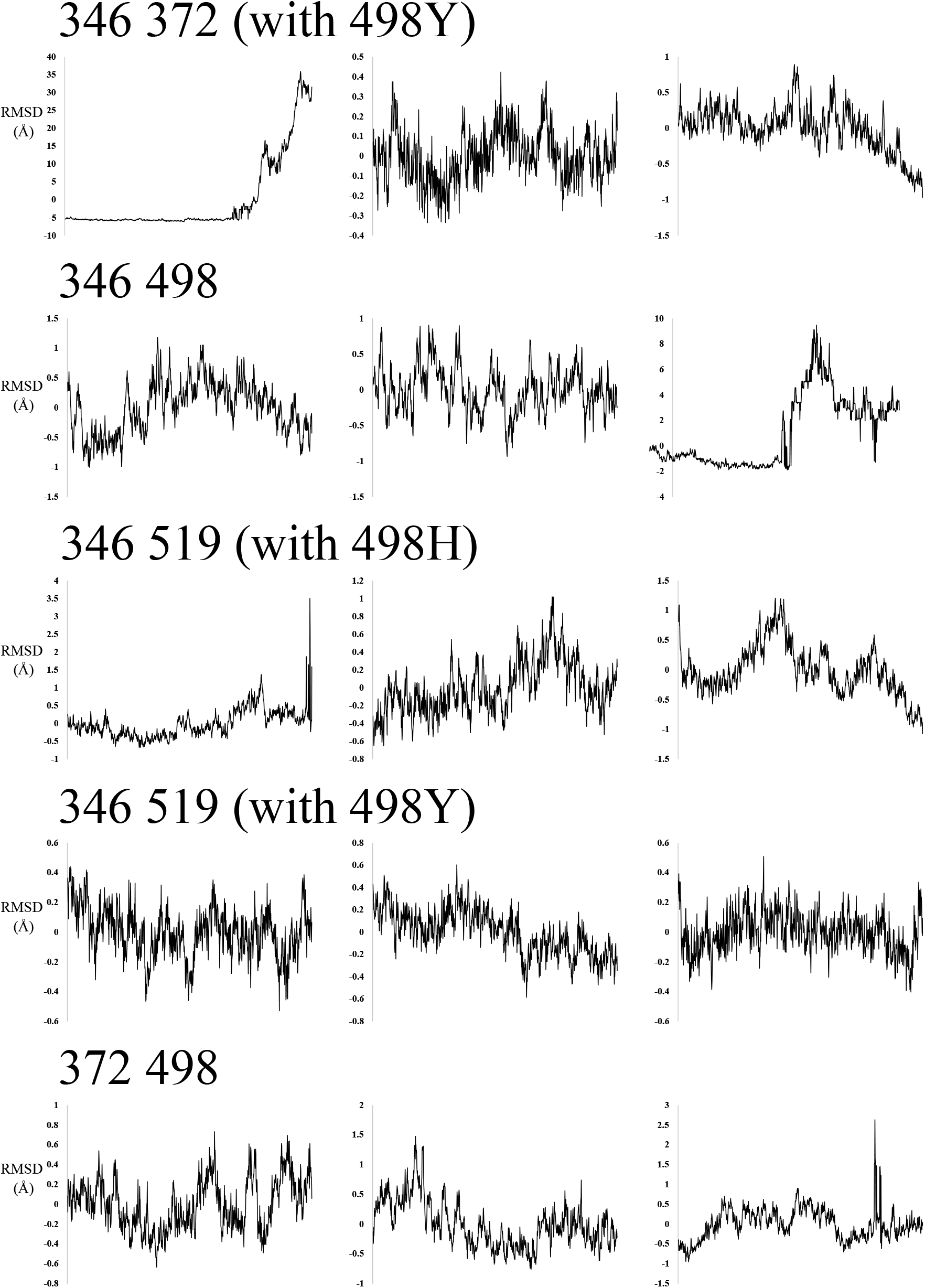
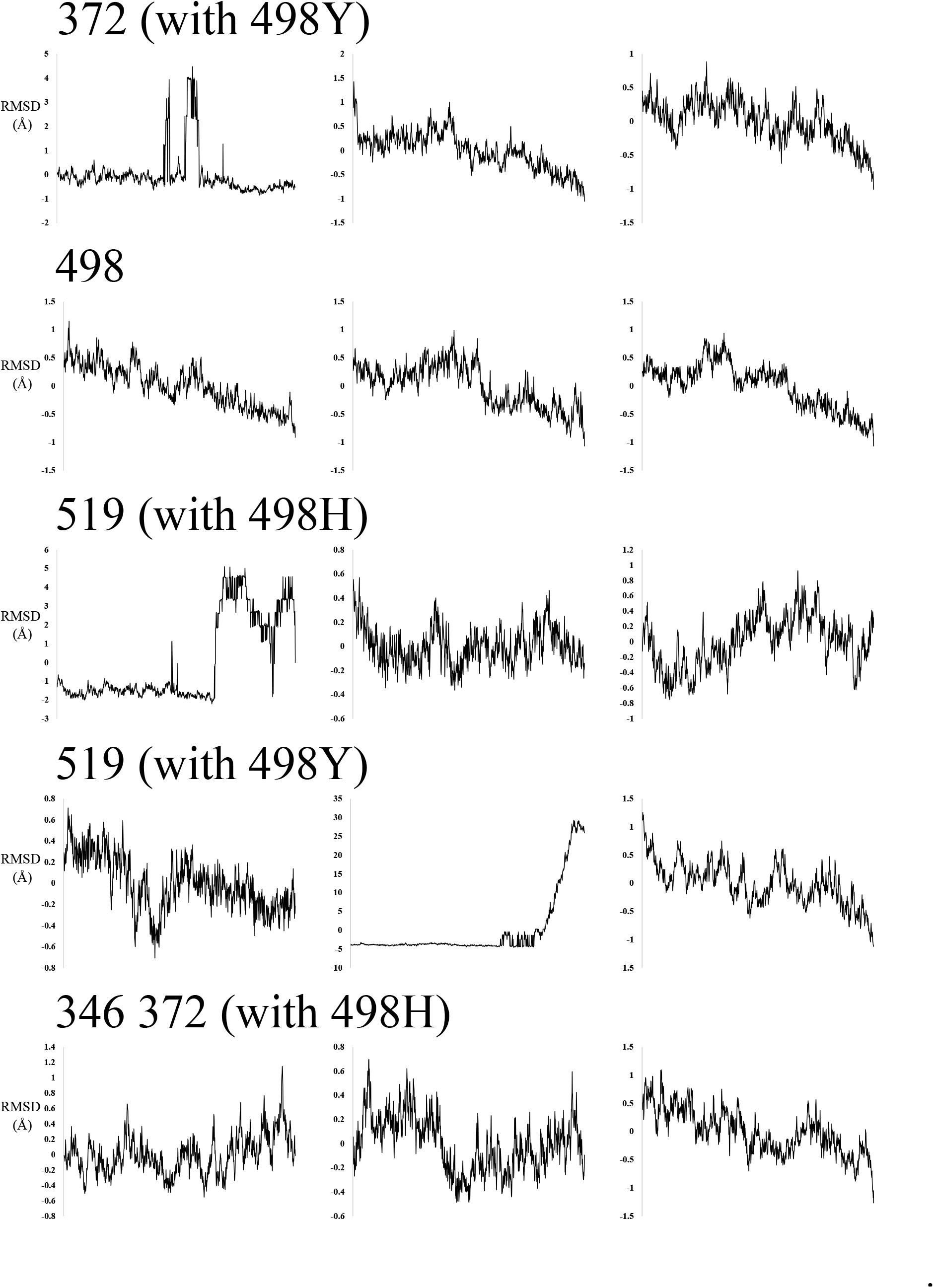
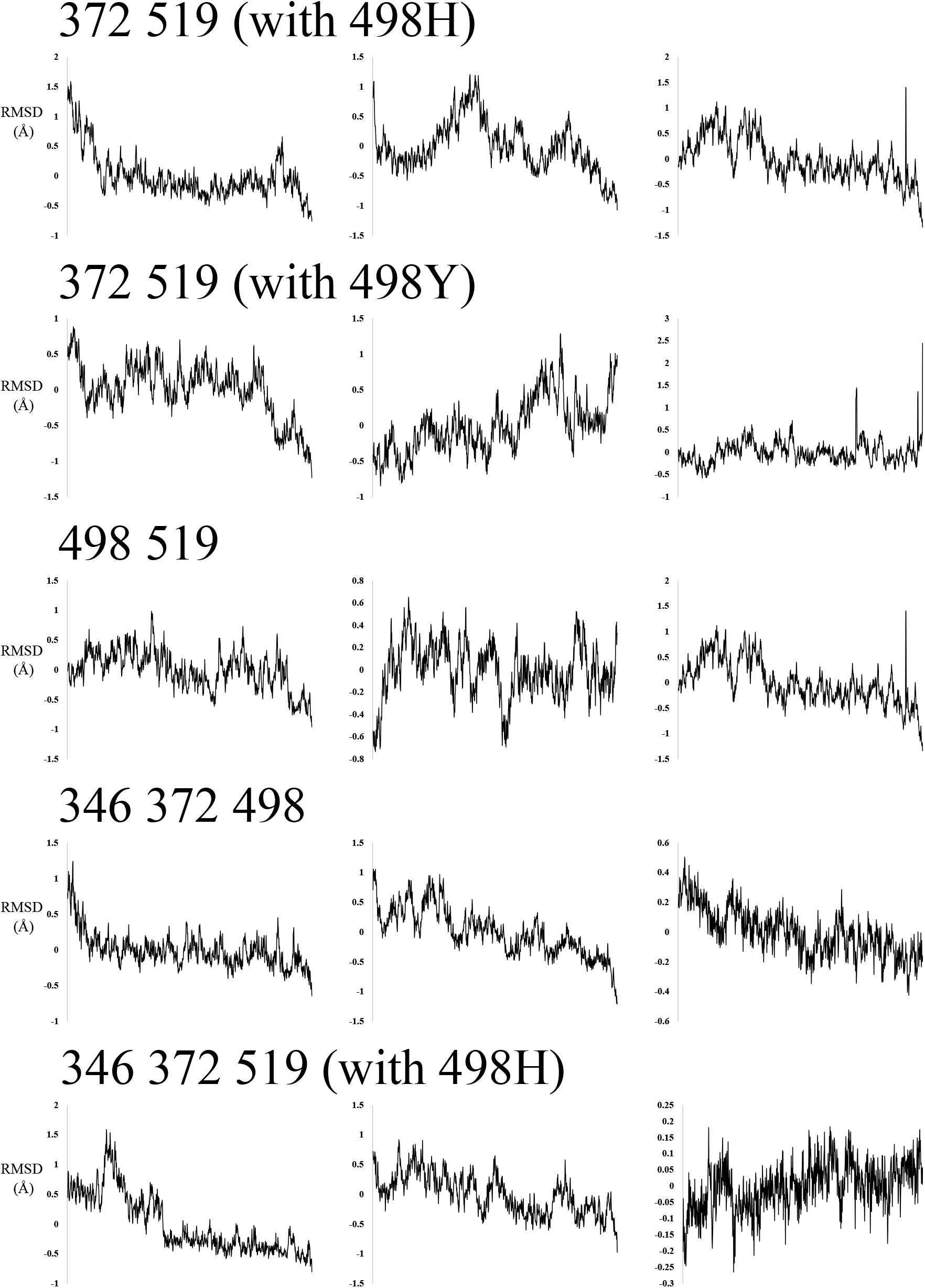
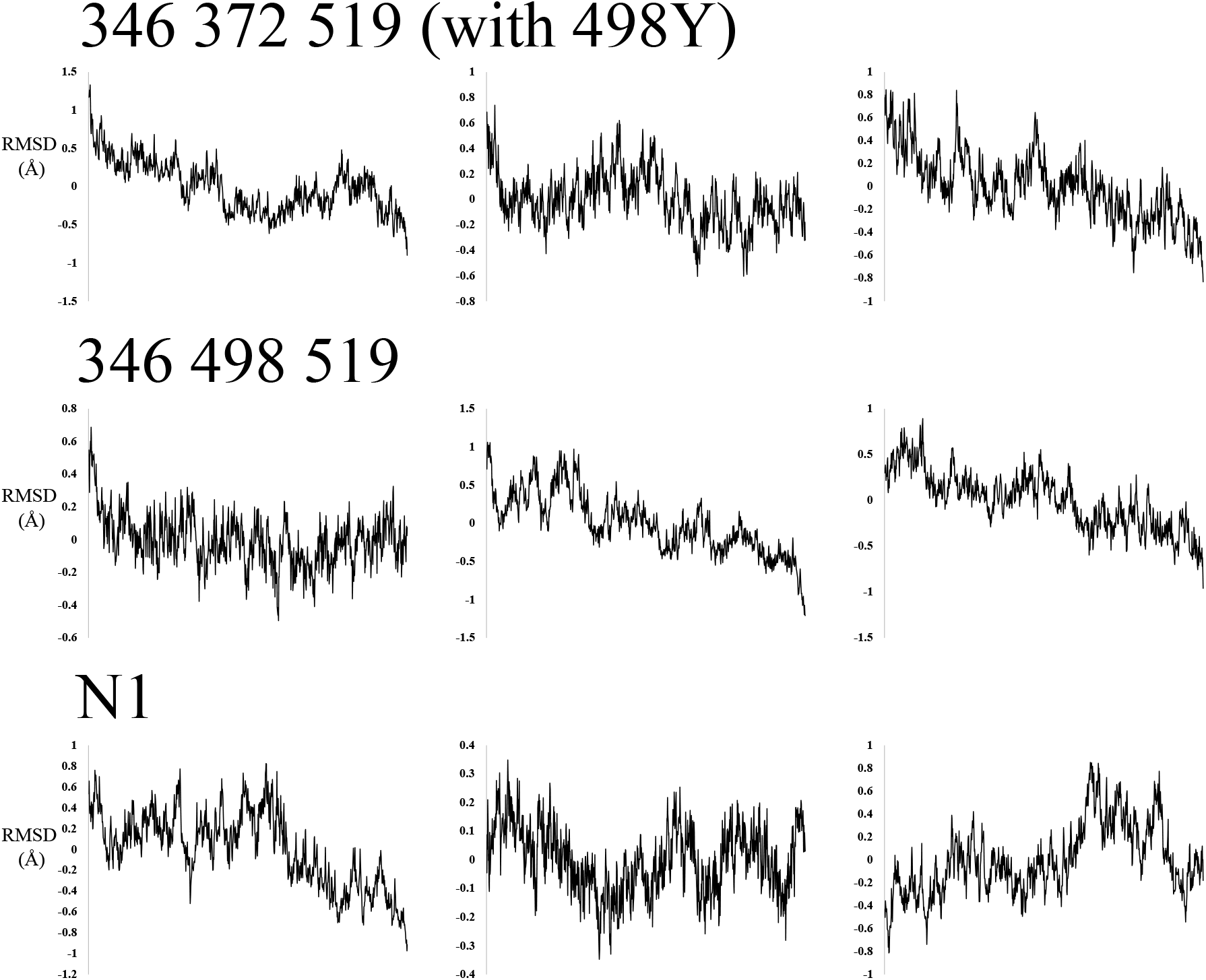
RMSD of simulation data used for energy calculations. Root-mean-square deviation (RMSD) is shown for simulation window that was used to calculate complex binding energy. Note that the Spike-RBD maintains a consistent stable configuration at the interface with hACE2, suggesting our energy calculations can be safely compared across simulations and that higher random stochasticity should not be a confounding factor.

**Supplementary Figure 3:**
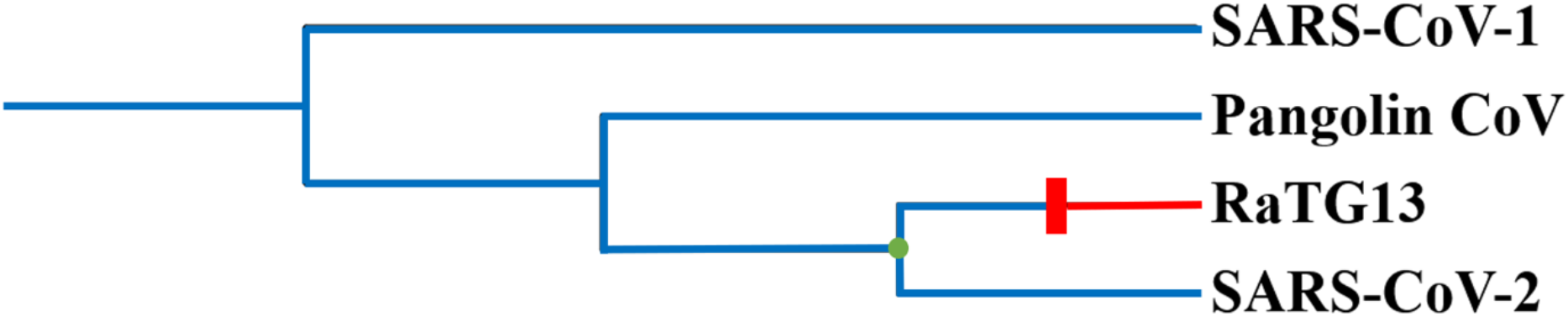
Maximum Parsimony Reconstruction of hACE2 binding by Spike-RBD. While weak evidence on its own, a reconstruction by maximum parsimony of Spike-RBD ability to bind hACE2 for the closest relatives of SARS-CoV-2 is consistent with our ancestral reconstruction here in suggesting that the common ancestor with RaTG13 possessed high binding affinity for hACE2.

**Supplemental Table 1:** Statistical confidence of ancestral sequence reconstructions for positions that vary between N0 and N1. Ancestral sequence reconstruction was assessed via computed posterior probability for each reconstructed state at each position in the sequence. The posterior probability for each reconstructed state at the four key positions that vary between N0 and N1 is shown, as calculated by two independent software packages (FastML and GRASP).

| FastMLreconstructions |  |  |  |  | GRASP reconstructions |  |  |  |
| --- | --- | --- | --- | --- | --- | --- | --- | --- |
| Position | N0 | Confidence | N1 | Confidence | N0 | Confidence | N1 | Confidence |
| 346 | T | 0.94 | R | 1 | T | 0.99 | R | 1 |
| 372 | T | 0.97 | A | 1 | T | 0.91 | A | 1 |
| 498 | H/Y | 0.3/0.61 | Q | 1 | H/Y | 0.48/0.40 | Q | 1 |
| 519 | N | 0.98 | H | 1 | N | 0.94 | H | 1 |

**Supplemental Table 2:** Sources of all SARS-CoV-2 genome sequences. Each genome sequence analyzed from SARS-CoV-2 infection cases are detailed according to geographic region of origin, as well as potentially relevant patient meta-data.

| Country | Number of Sequences |  |  |  | Average Age of Sequenced Patients |  |  |  |
| --- | --- | --- | --- | --- | --- | --- | --- | --- |
|  | Male | Female | Unknown | All | Male | Female | Unknown | All |
| Algeria | 1 | 1 | 0 | 2 | 97.0 | 28.0 | N/A | 62.5 |
| Australia | 11 | 15 | 1 | 27 | 38.5 | 36.0 | N/A | 37.1 |
| Belgium | 14 | 9 | 0 | 23 | 40.6 | 54.2 | N/A | 45.9 |
| Brazil | 5 | 3 | 0 | 8 | 37.0 | 29.0 | N/A | 34.0 |
| Cambodia | 1 | 0 | 0 | 1 | 60.0 | N/A | N/A | 60.0 |
| Canada | 10 | 7 | 0 | 17 | 64.5 | 54.7 | N/A | 60.5 |
| Chile | 4 | 3 | 0 | 7 | 30.5 | 43.0 | N/A | 35.9 |
| China | 29 | 7 | 7 | 43 | 48.9 | 47.7 | N/A | 48.7 |
| Congo | 3 | 2 | 0 | 5 | 45.0 | 28.5 | N/A | 38.4 |
| Czech Rep | 1 | 0 | 0 | 1 | 44.0 | N/A | N/A | 44.0 |
| Denmark | 4 | 1 | 0 | 5 | 40.5 | 21.0 | N/A | 36.6 |
| Finland | 4 | 4 | 0 | 8 | 67.3 | 38.3 | N/A | 52.8 |
| France | 17 | 11 | 3 | 31 | 62.0 | 63.2 | 56.0 | 61.8 |
| Georgia | 8 | 2 | 0 | 10 | 39.9 | 40.0 | N/A | 39.9 |
| Germany | 1 | 1 | 9 | 11 | N/A | 38.0 | N/A | 38.0 |
| Greece | 0 | 0 | 1 | 1 | N/A | N/A | N/A | N/A |
| Hong Kong | 10 | 11 | 0 | 21 | 57.3 | 51.3 | N/A | 54.1 |
| Hungary | 2 | 1 | 0 | 3 | 26.5 | 26.0 | N/A | 26.3 |
| Iceland | 5 | 13 | 0 | 18 | 44.4 | 46.8 | N/A | 46.1 |
| India | 1 | 0 | 0 | 1 | 23.0 | N/A | N/A | 23.0 |
| Ireland | 2 | 2 | 0 | 4 | 54.0 | 25.5 | N/A | 39.8 |
| Italy | 9 | 2 | 0 | 11 | 56.0 | 72.5 | N/A | 59.0 |
| Japan | 1 | 2 | 8 | 11 | 60.0 | 62.0 | N/A | 61.3 |
| Kuwait | 0 | 0 | 1 | 1 | N/A | N/A | N/A | N/A |
| Luxembourg | 0 | 1 | 1 | 2 | N/A | 32.0 | N/A | 32.0 |
| Malaysia | 1 | 2 | 0 | 3 | 11.0 | 40.0 | N/A | 30.3 |
| Mexico | 1 | 0 | 0 | 1 | 35.0 | N/A | N/A | 35.0 |
| Nepal | 1 | 0 | 0 | 1 | 32.0 | N/A | N/A | 32.0 |
| Netherlands | 0 | 0 | 14 | 14 | N/A | N/A | N/A | N/A |
| New Zealand | 1 | 2 | 0 | 3 | N/A | 60.0 | N/A | 60.0 |
| Norway | 0 | 0 | 6 | 6 | N/A | N/A | N/A | N/A |
| Pakistan | 0 | 1 | 0 | 1 | N/A | 40.0 | N/A | 40.0 |
| Panama | 0 | 1 | 0 | 1 | N/A | 40.0 | N/A | 40.0 |
| Peru | 0 | 1 | 0 | 1 | N/A | 61.0 | N/A | 61.0 |
| Poland | 1 | 0 | 0 | 1 | 66.0 | N/A | N/A | 66.0 |
| Portugal | 10 | 4 | 1 | 15 | 32.3 | 34.5 | N/A | 33.1 |
| Russia | 0 | 1 | 0 | 1 | N/A | 30.0 | N/A | 30.0 |
| Saudi Arabia | 1 | 1 | 0 | 2 | 68.0 | 67.0 | N/A | 67.5 |
| Senegal | 5 | 2 | 0 | 7 | 44.4 | 55.0 | N/A | 47.4 |
| Singapore | 8 | 4 | 0 | 12 | 45.7 | 44.5 | N/A | 45.2 |
| Slovakia | 1 | 1 | 0 | 2 | 26.0 | 59.0 | N/A | 42.5 |
| South Africa | 1 | 0 | 0 | 1 | 38.0 | N/A | N/A | 38.0 |
| South Korea | 4 | 0 | 2 | 6 | 46.3 | N/A | N/A | 46.3 |
| Spain | 13 | 3 | 0 | 16 | 61.5 | 47.7 | N/A | 58.9 |
| Sweden | 0 | 0 | 1 | 1 | N/A | N/A | N/A | N/A |
| Switzerland | 2 | 0 | 9 | 11 | 49.0 | N/A | 50.2 | 50.0 |
| Taiwan | 4 | 10 | 0 | 14 | 52.0 | 56.2 | N/A | 55.0 |
| Thailand | 0 | 1 | 0 | 1 | N/A | 74.0 | N/A | 74.0 |
| UK | 17 | 12 | 0 | 29 | 55.2 | 50.6 | N/A | 53.4 |
| USA | 7 | 8 | 38 | 53 | 38.7 | 52.0 | N/A | 45.8 |
| Vietnam | 1 | 2 | 1 | 4 | 50.0 | 57.0 | N/A | 54.7 |
| All Countries | 222 | 154 | 103 | 479 | 49.1 | 47.8 | 51.7 | 48.7 |

**Supplementary Table 3.** Results of an equivalence test for means. Equivalence test compares z-score standardized molecular dynamics data to z-score standardized *in vitro* data. Test illustrates that the molecular dynamics simulation reflects *in vitro* SARS-CoV-2 binding energies (p = 0.042).

| Equivalence Test for Means | | $\alpha = 0.05$ |
| --- | --- | --- |
| Equal Sample Sizes |  |  |
|  | *MD* | *Starr et al.* |
| Mean | 1.7901 | 0.5676 |
| Variance | 0.5824 | 1.25 |
| Observations | 5 | 5 |
| Hypothesized Mean Difference | 0 |  |
| df | 7 |  |
| t Stat | 2.019 | -2.019 |
| p1 (MD) and p2 (Starr et al.) | 0.042 | 0.042 |
| T Critical one-tail | 1.895 |  |
| Means are Equivalent because p1 & p2 < 0.05 |  |  |

